# The ZNF512B-NuRD complex co-ordinates a neural-specific gene expression program

**DOI:** 10.1101/2025.06.23.660971

**Authors:** Habib Francis, Mehdi Sharifi Tabar, Chirag Parsania, Yue Feng, Caroline Giardina, Omar Ibrahim, Krishna P Singh, Shailendra K Gupta, Ulf Schmitz, John EJ Rasko, Charles G Bailey

## Abstract

The nucleosome remodelling and deacetylase complex (NuRD) plays a key role in chromatin regulation and a wide range of biological processes including development, haemopoiesis, immunity and neurogenesis. Its interaction with tissue-enriched and sequence-specific transcription factors (TFs) leads to distinct functional outputs in the given tissue by targeting a specific set of genes. However, how NuRD dynamically and specifically regulates gene expression in a tissue-specific manner is poorly understood. Here, we refine an N-terminal specific NuRD interaction motif which enables direct engagement with many transcriptional regulatory proteins. Using a series of structural modelling and biochemical techniques, we show that ZNF512B, a poorly characterised neuronal-expressed zinc finger protein, directly binds to the RBBP4 subunit of the NuRD complex. Subsequent knockdown of ZNF512B results in the downregulation of several neural-related molecular pathways suggesting that ZNF512B may play a regulatory role during neurogenesis. We also show that in NTERA-2 neural cells, the expression of ZNF512B is necessary for cell growth and survival, and is markedly enhanced during neural progenitor cell (NPC) differentiation. In summary, our data suggest that ZNF512B might regulate neural-specific transcriptional programs via engagement with the NuRD complex.

## Introduction

The nucleosome remodelling and deacetylase (NuRD) complex is an evolutionarily conserved chromatin remodelling complex in metazoa^1,2^. The complex consists of seven core subunits, many with paralogs, including CHD3, CHD4 and CHD5, HDAC1 and HDAC2, MBD2 and MBD3, RBBP4 and RBBP7, GATAD2A and GATAD2B, MTA1, MTA2 and MTA3, and CDK2AP1. CHD and HDAC subunits constitute the enzymatic components responsible for nucleosome sliding and acetyl mark removal from lysine residues on histone tails, respectively. The remaining subunits are mainly responsible for DNA, histone and protein-protein interactions^3–5^. It is notable that CHD, HDAC, RBBP, and CDK2AP1 subunits are also found in other protein complexes involved in gene regulation^6–8^. NuRD plays a role in various molecular and cellular pathways during development and disease, partly through paralog switching. For example, MBD2-NuRD but not MBD3-NuRD is the major suppressor of fetal hemoglobin gene in human^9–11^. GATAD2A-but not GATAD2B-NuRD inhibits induced pluripotent stem cell formation from somatic cells^12^. Besides paralog switching-associated functional outputs, interaction of NuRD with a range of transcription factors (TFs) can lead to distinct functional outcomes^13,14^.

Lineage-specific TF assembly with the NuRD complex generates specialized transcriptional machinery which can localize at specific genomic loci across the genome. TFs show dynamic expression patterns during development or cellular differentiation, leading to dynamic and cell-type-specific binding of NuRD to promoters and enhancer regions^15^. For example, in the tumour microenvironment, the PAC-1-NuRD complex specifically regulates the gene expression program of exhausted T cells^13^. In addition, IKAROS interaction with the NuRD complex regulates hematopoietic stem cell growth and differentiation and immune cell fate determination^16^. In the context of neurogenesis, both *in vitro* and *in vivo* studies have revealed that the NuRD complex regulates a distinct gene expression program in neuronal cells^17–20^. This has been shown to be partly through deposition of the H2A.Z histone variant at the promoter of brain-specific genes by the NuRD complex^20^. However, how NuRD is recruited to specific promoter or enhancer regions in specific populations of cells within the brain remains elusive. Identification and characterisation of neuronal-specific TF-NuRD complexes will help delineate the role of transcriptional programs during neurogenesis.

Zinc finger (ZF) proteins, comprising the largest family of TFs, modulate diverse biological processes and are mainly involved in transcriptional gene regulation^21,22^. For a significant proportion of ZF proteins (∼50%), little is known about their biological function. Interestingly, a number of genetic variants in ZF proteins have been documented in patients with neurodegenerative diseases^23–25^. For one such protein, ZNF512B, single nucleotide polymorphisms in multiple regions of the *ZNF512B* locus are associated with Amyotrophic Lateral Sclerosis (ALS) disease^26–29^. ALS is a severe neurodegenerative disease affecting motor neurons and neural progenitor cell (NPC) function. However, the molecular and biological function of ZNF512B, a multi-ZF-containing protein, remains elusive. A clearer understanding of ZNF512B function will help delineate its contribution to neurogenesis in health and disease.

In this study, we provide strong correlative evidence for ZNF512B as a NuRD interacting protein from protein interaction databases and Alphafold structural prediction. Then, using an proteomics approach we confirm that ZNF512B physically interacts with the RBBP4 subunit of the NuRD complex via a conserved NuRD interaction motif (NIM). We further show that depletion of ZNF512B leads to the downregulation of many genes with a defined function in various neuronal pathways. Finally, using an NTERA-2 cell differentiation model, we demonstrate that ZNF512B expression is markedly enhanced in NPCs as compared to non-NPC cells.

## Results

### Refinement of the NuRD interaction motif

The NuRD interaction motif (NIM) previously defined as being necessary for engagement with the NuRD complex is RRKxxxP. A recent study proposed an alternative internal NIM, RKxxxPxK^30^. To refine this further we collated all human proteins that contained putative NIMs using Scansite and examined their overlap (**Figure 1A, Supplementary Table 1**). We cross-matched these 317 proteins with the BioGRID protein-protein interaction database to determine if independent evidence for interaction with NuRD subunits existed. Approximately 34% (45 out of 178) of RRKxxxP-containing proteins interacted with NuRD complex subunits, whereas only 21% (33 out of 156) of RKxxxPxK-containing proteins did (**Figure 1B**). Their respective sequence logos could not be further refined based on all sequence evidence (**Figure 1C**, *top* & *upper middle panel*). However, based on the 45 NuRD-interacting proteins containing RRKxxxP identified in BioGRID, we observed an emergent consensus ‘RRKQxxP’ (**Figure 1C**, *lower middle panel*). Therefore, refining our Scansite search to RRKQxxP, we identified 23 proteins, with 19 (83%) exhibiting prior evidence of NuRD subunit interaction (**Figure 1B**), however the NIM could not be further refined (**Figure 1C**, *bottom panel*). We then examined the relative position of each NIM within all proteins in each subset. The RRKxxxP motif was distributed mostly in the first half of proteins, whereas, the RRKQxxP motif, was predominantly enriched at the N-terminus. The N- and C-termini were devoid of the RKxxxPxK-based NIM, consistent with its description as an internal NIM^30^ (**Figure 1D**). Using the DAVID bioinformatics resource, proteins experimentally validated to associate with RRKQxxP-containing proteins (BioGRID) highlighted that retinoblastoma binding protein 4 (RBBP4) was the predominant protein enriched and also a subunit of the NuRD complex (**Figure 1E**). Not surprisingly, transcriptional regulation was the major biological function enriched (**Figure 1F**). Consistent with this gene regulatory function, we examined the subcellular localisation of all NIM-containing proteins and showed that the majority of RRKQxxP-containing proteins were nuclear-localised (**Supplementary Figure 1A**). Furthermore, the majority are proteins not essential for cell viability (**Supplementary Figure 1B**). Interestingly, over 85% of proteins were associated with a known genetic disease (**Supplementary Figure 1C**). Examining the known functions of all RRKQxxP-containing proteins identified that most of these transcriptional regulatory proteins exhibit specialised functions when engaged with the NuRD complex (**Table 1**). Many N-terminal NIM-containing proteins have been well-characterised, however the internal RRKQxxP-containing protein, ZNF512B, is less functionally characterised, and thus the focus of this study.

**Figure 1.**
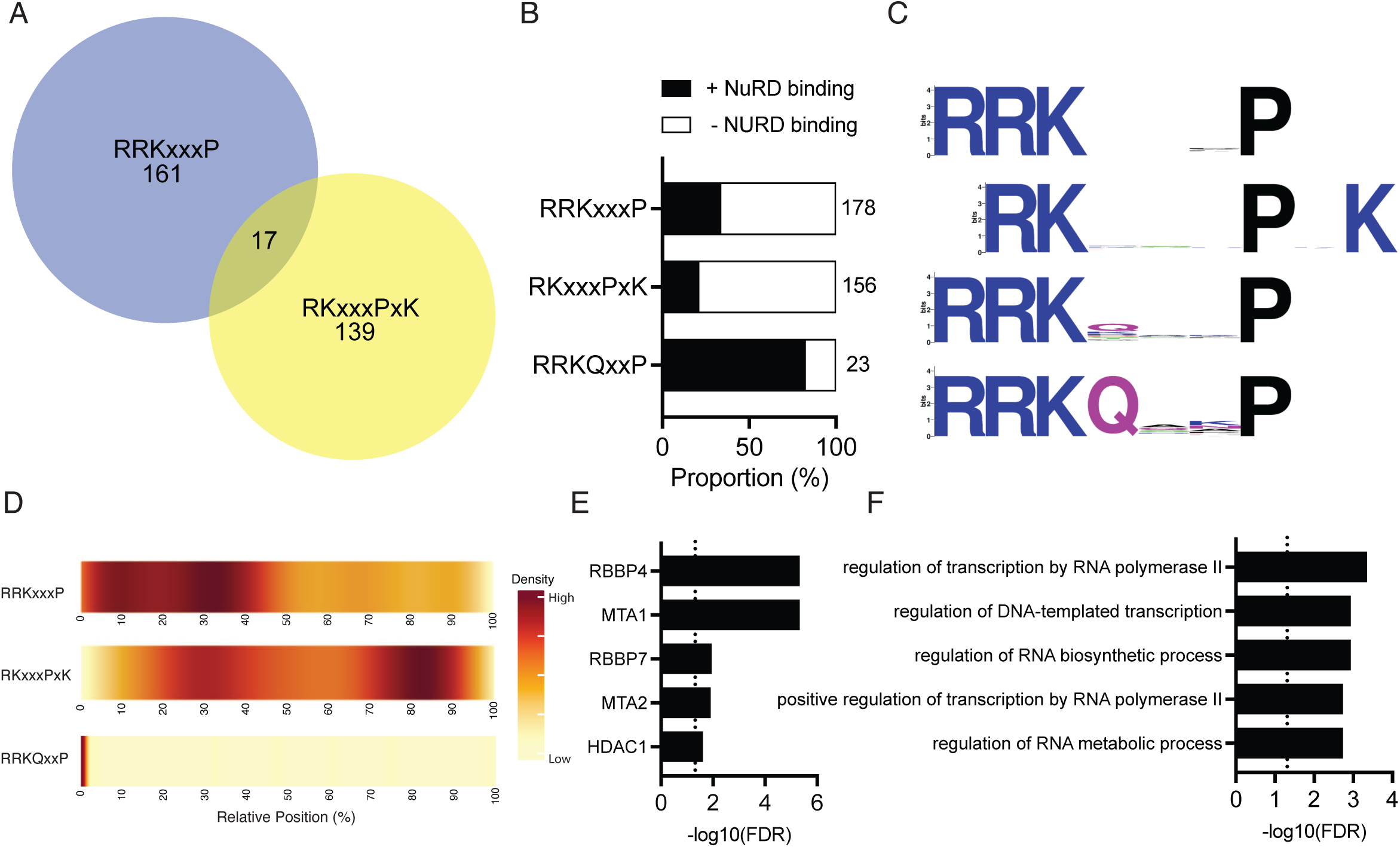
The RRKQxxP motif is enriched in the N-terminus of transcriptional regulatory proteins interacting with NuRD. **(A)** Occurrence of NIM sequences in human proteins; numbers for each NIM are indicated. **(B)** Proportion of NIM-containing proteins with evidence of protein interaction with NuRD subunits. The total number of human proteins are shown; data was manually curated from BioGRID. **(C)** Sequence logos generated from multiple sequence alignments of NIMs. **(D)** The relative location of the NIM in all NIM-containing proteins with the colour scale representing density. **(E-F)** RRKQxxP NIM-containing proteins were analysed for: **(E)** the most common interacting proteins; and **(F)** enrichment of biological processes; using DAVID.

**Table 1.** Summary and biological role of proteins identified with the RRKQxxP NIM, many of which have previously been shown to functionally interact with the NuRD complex.

| Function | Protein | Role with NuRD |
| --- | --- | --- |
| Transcription regulator | FOG1, FOG2 | Megakaryocytic & erythroid development; cardiogenesis |
| Transcription factor | BCL11A, BCL11B | Lymphopoietic development |
| Transcription factor | SALL1, SALL2, SALL3, SALL4 | Organogenesis; stem cell renewal |
| Transcription factor | TSHZ1, TSHZ2, TSHZ3 | Developmental processes |
| Transcription factor | ZEB1, ZEB2 | Regulation of epithelial-mesenchymal transition |
| Transcription factor | ZNF827 | Telomere maintenance |
| Transcription factor | ZNF521 | Neural differentiation |
| Transcription factor | ZNF512B, ZNF513, ZNF821 | Transcriptional regulation? |
| Signal transduction | DVL2 | Wnt signalling |
| Miscellaneous | CND2, KRI1, LZTR1, MYCT1 | N.D. |

### Structural modelling confirms ZNF512B-RBBP4 interaction

To independently verify the interaction likelihood of ZNF512B with NuRD subunit RBBP4, we performed pairwise interaction analysis of ZNF512B with all 15 subunits of the NuRD complex using Alphafold-Multimer. This confirmed that ZNF512B had a high probability of interaction with RBBP4 or RBBP7 as determined by their interface predicted template modelling (iPTM) score, measuring 0.79 and 0.8 respectively (**Figure 2A**). Examining the ZNF512B-RBBP4 interaction in more detail by paired residue analysis using Alphafold revealed high confidence interactions which are depicted in a heatmap with low prediction aligned error (PAE) scores (**Figure 2B**). This analysis confirmed an interaction interface within ZNF512B, which overlapped with residues 421-427 where the ZNF512B NIM RRKQKTP is located (**Figure 2B**).

**Figure 2.**
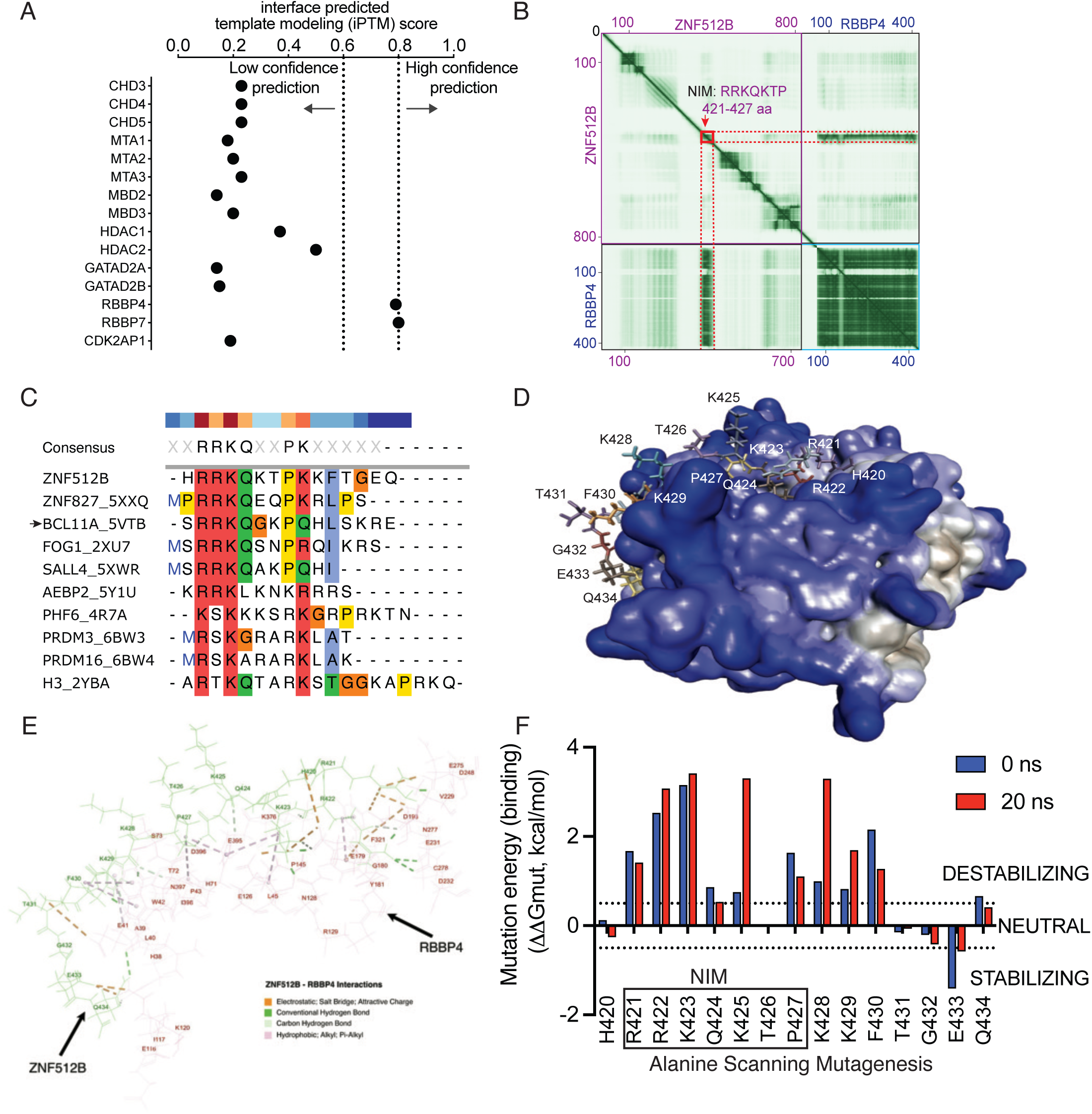
Alphafold and structural modelling predict ZNF512B interaction with RBBP4. **(A)** Alphafold-Multimer analysis of ZNF512B binary interactions with individual NURD complex subunits. Interface predicted template modelling (iPTM) thresholds of 0.6 and 0.8 indicate low and high confidence predictions respectively. **(B)** Heatmap representing the predicted aligned error (PAE) scores between all pairs of residues of ZNF512B and RBBP4. The PAE scores are depicted across all residue pairs, with dark green indicating low PAE (high confidence in relative positioning) and lighter shades reflecting lower prediction confidence. Red box indicates the predicted interaction interface in ZNF512B containing the NIM (residues 421-427) that forms with RBBP4. **(C)** Sequence alignment of all NIM-containing peptides used in published RBBP4 structural studies. The protein name and PDB accession name are indicated; arrow indicates BCL11A sequence and structure used in this study. **(D)** A 3D molecular model of ZNF512B peptide (residues 420-434) with RBBP4, obtained by substituting in ZNF512B residues for the BCL11A peptide bound to RBBP4. **(E)** All bond types occurring between ZNF512B and RBBP4 over a 20 ns molecular dynamics simulation run are depicted in the stick model. **(F)** Conformational stability of the ZNF512B-RBBP4 model after each individual residue was mutated to alanine and molecular dynamics simulations were performed. The mutation energy (binding) was measured at the start and end of the 20 ns run. The thresholds for decreased binding affinity (ΔΔGmut >0.5 = destabilising) and increased binding affinity (ΔΔGmut <0.5 = stabilising) are indicated.

To generate a ZNF512B-RBBP4 3D structural model, we required suitable existing RBBP4-containing complexes which exhibited a similar binding mode. From the Protein Data Bank, 9 RBBP4 structures were identified along with peptides representing NIM-containing proteins (**Figure 2C**). BCL11A NIM-containing peptide (residues 2-16) bound to RBBP4^31^ was chosen as the reference model due to its similarity to ZNF512B NIM. We extracted the structure, performed structure optimisation and energy minimisation, pinpointed the RBBP4 docking interface for the BCL11A peptide, and then corresponding residues in ZNF512B (420-434) were substituted in using molecular modelling. The resulting 3D model shows the 15 residue ZNF512B peptide binding on the acidic surface and within an uncharged pocket of RBBP4 (**Figure 2D**). All bonds occurring between ZNF512B and RBBP4 over a 20 ns run were visualized and documented (**Supplementary Table 2 & 3**). Our analysis indicated that R421, R422, K423, Q424 and P427 are the primary residues within the NIM motif involved in contact within the interaction interface (**Figure 2E**). These include conventional and carbon hydrogen bonds, electrostatic and hydrophobic bonds. Close-up views of individual residues from the ZNF512B NIM motif are shown (**Supplementary Figure 2**). Other auxiliary residues in ZNF512B including H420, K429, F430, E433 and Q434 exhibit a range of bond types which would stabilise the interaction (**Figure 2E**). The sidechains of K425, T426 and K428 residues are projected away from RBBP4, consistent with a previous report^30^.

We next predicted the importance of individual residues for the ZNF512B-RBBP4 interaction by generating mutant models featuring individual alanine substitutions of all 15 ZNF512B residues. Molecular dynamics simulations were performed for 20 ns to examine the conformational stability of the ZNF512B-RBBP4 mutant models. Mutation of R421, R422, K423, P427 and F430 residues to alanines all individually destabilised the ZNF512B-RBBP4 interaction before and after the simulation run (**Figure 2F**). Lysines at positions 425, 428 and 429 were further destabilised by mutation after 20 ns simulation (**Figure 2F**). As expected, mutations in the NIM had the highest destabilising effects on ZNF512B-RBBP4 interaction which was reflected in the number of bonds lost upon mutation when compared to WT (**Supplementary Table 2 & 3**). These data provides additional supportive and independent evidence for the interaction of ZNF512B with RBBP4 via the NIM motif.

### ZNF512B interacts with the NuRD complex via RBBP4

One of the features of ZF proteins is their interaction with multi-subunit protein complexes which influence gene expression programs^32^. To map ZNF512B protein interaction partners, we performed ZNF512B pulldown followed by mass spectrometry analysis (PD-MS). A previous study used a similar approach with an ZNF512B-GFP fusion featuring a large C-terminal ∼27 kDa eGFP tag^30^, which may affect protein complexes. To avoid this, we expressed an N-terminal (∼2 kDa) haemagglutinin epitope (HA)-tagged full-length ZNF512B and empty vector containing HA-tag only as control (CTRL) in Expi293F cells. Using ZNF512B as bait, PD-MS data analysis revealed a marked enrichment of most canonical subunits of the NuRD complex compared to the control (**Figure 3A**). To further corroborate our finding, we conducted pairwise co-immunoprecipitation (Co-IP) experiments of HA-tagged ZNF512B with FLAG-tagged core subunits of the NuRD complex: CHD4-C (C-terminus of CHD4 protein that interacts with the NuRD complex^33^), MBD3, MTA2, RBBP4, and GATAD2B. Plasmids were co-expressed in Expi293F cells for 72 h, and cell lysates were mixed with anti-HA beads to capture ZNF512B and interacting NuRD components. Western blot analysis of the eluates revealed equivalent enrichment of ZNF512B at 120 kDa using a HA antibody in all samples, and varying co-purification of all NuRD subunits as revealed with a FLAG antibody using a standard washing buffer containing 200 mM NaCl (**Figure 3B**). RBBP4 was more enriched compared to other NuRD subunits (CHD4, MBD3, MTA2 and GATA2B), where only weak interactions were observed. These weak interactions are probably mediated by indirect bridging of endogenous NuRD subunits, an issue we had previously described^2,34^. Notably, all experiments were performed in the presence of the nuclease benzonase to minimise DNA- or RNA-mediated interactions.

**Figure 3.**
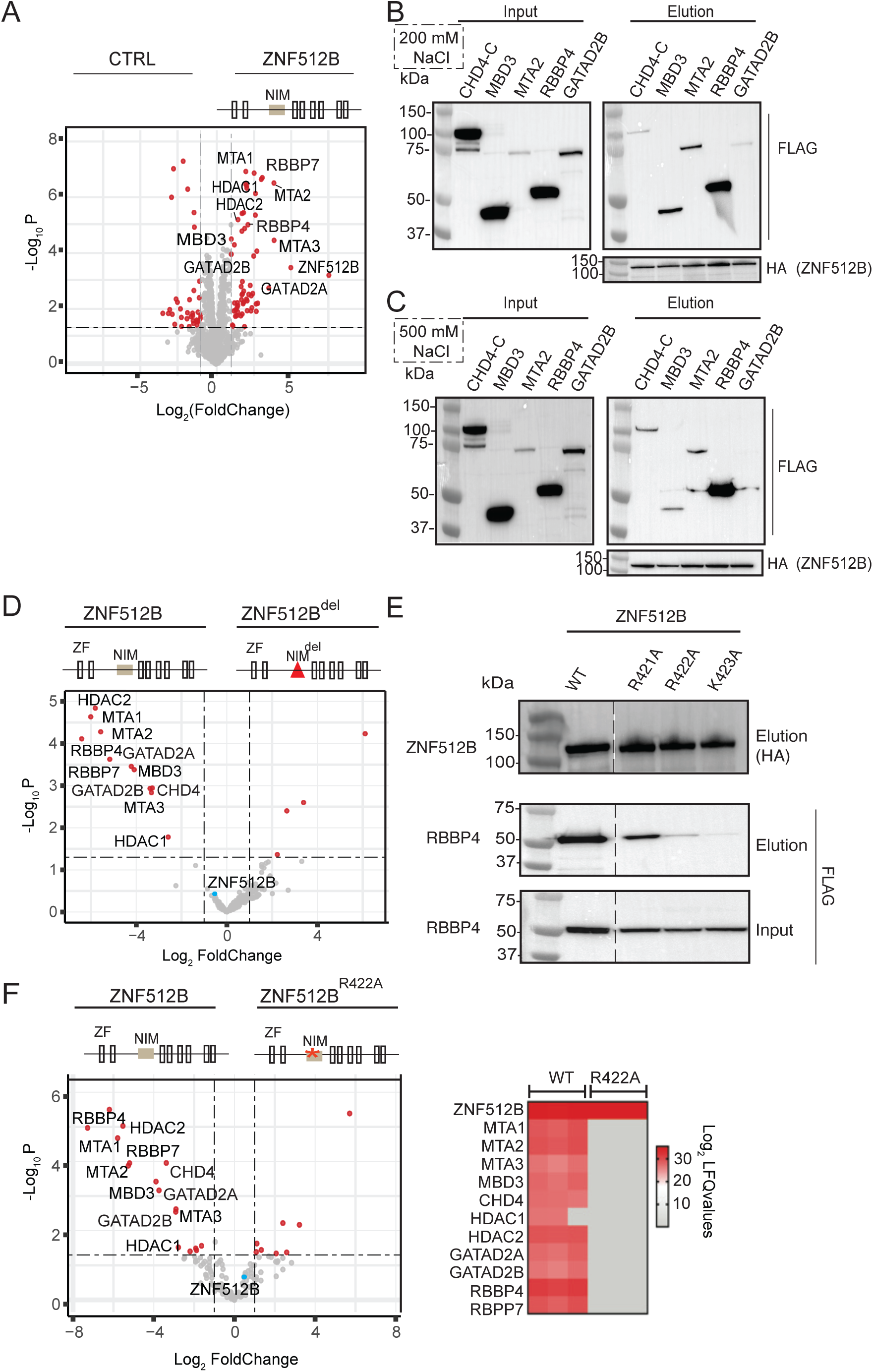
ZNF512B interacts with the RBBP4 subunit of the NuRD complex via the internal NIM. **(A)** Volcano plot represents the affinity purification of HA-tagged ZNF512B and its binding partners in Expi293F cells. Enrichment of most canonical subunits of the NuRD complex was observed as compared to the CTRL (HA-only) samples (left). Significantly enriched proteins are indicated with red dots and non-significant proteins are indicated with grey dots. **(B-C)** CoIP of HA-ZNF512B (bait) with FLAG-tagged NuRD subunits (prey) followed by Western blot analysis. FLAG-tagged CHD4-C, MBD3, MTA2, and GATAD2B were co-expressed individually with HA-ZNF512B for 72 h, and cell lysates were mixed with HA-beads to capture ZNF512B and the associated NuRD subunit. Beads were washed in buffer containing 200 mM **(B)** and 500 mM NaCl **(C)**. (**D)** Volcano plot demonstrating the loss of NuRD subunits enrichment when HA-ZNF512B^del^ (right panel) was used as a bait as compared to HA-ZNF512B (left panel). As expected, ZNF512B (blue dot) is not enriched. Schematics of the ZNF512B constructs are indicated on top of the volcano plots. **(E)** Western blot showing co-purification of FLAG-RBBP4 (prey) with HA-ZNF512B (bait) protein carrying different NIM mutations. The input blot was probed with anti-HA and the elution was developed with both anti-HA and anti-FLAG antibodies. Expected size for ZNF512B is 97 kDa, however; it migrates at 130 kDa possibly due to post-translational modifications affecting its size. Dashed line indicates that the blot was initially cut, and HA-ZNF512B wildtype (WT)- and mutant-containing membranes were joined for visualization purposes. **(F)** *Left panel*: Volcano plot comparing the PD-MS protein interaction profile in ZNF512B^R422A^ pulldown (right) versus ZNF512B or WT (left). *Right panel*: Heatmap of LFQ intensities for all NuRD subunits after PD-MS. Data represents the mean of three tested replicates.

To further refine the potential direct binding partner of ZNF512B, we repeated the same Co-IP with more stringent washing conditions (i.e. 500 mM NaCl-containing buffer). Again, RBBP4 was highly enriched as compared to other NuRD subunits (**Figure 3C**) suggesting that RBBP4 was a direct binding partner of ZNF512B. As our sequence analysis, structural modelling and a recent study^30^ provided evidence that this interaction occurs through the NIM in ZNF512B (residues 421-427), we deleted the entire NIM (denoted as ZNF512B^del^) and performed PD-MS analysis along with ZNF512B as a positive control. Upon equivalent expression of both ZNF512B baits in the analysis, all NuRD subunits were significantly enriched with ZNF512B, whereas these interactions were completely interrupted when the NIM was deleted (**Figure 3D**). To further corroborate our PD-MS data, we also performed a Co-IP experiment followed by western blot analysis, where we individually mutated the ‘RRK’ residues in the NIM (amino acids 421-423) to alanines. RBBP4 was co-purified with ZNF512B but was reduced or lost when ZNF512B mutants were used as bait (**Figure 3E**). Depletion of all NuRD subunits with the ZNF512B R422A mutant (ZNF512B^R422A^) was further confirmed by PD-MS analysis, confirming that the NIM is essential for interaction of ZNF512B with the NuRD complex (**Figure 3F**).

### ZNF512B is evolutionarily conserved and intolerant to genetic loss-of-function and variation

ZNF512B is a 97 kDa protein with four tandem pairs of atypical (C2HC) and typical (C2H2) ZFs (**Figure 4A, Supplementary Figure 3A**), for which a DNA binding role was recently identified^30^. Apart from the interdispersed ZF pairs which are structured, ZNF512B is largely disordered (**Figure 4B**). The NIM is centrally located at residues 421-427 in a disordered region (**Figure 4A, B**), recently categorised as an internal NIM^30^. As little is known about a direct role of *ZNF512B* in human pathophysiology, we collated all available SNPs in the GnomAD database to examine the potential impact of normal genetic variation in *ZNF512B* (**Figure 4C**). There was a significant de-enrichment of missense SNPs occurring in the ZF domains of ZNF512B collectively (*p*<0.0003, **Table 2**). Also, significantly less missense SNPs were observed in ZFs 1, 2 & 4 than expected, suggesting these ZFs may be functionally important (all *p*<0.05, **Figure 4C**, **Table 2**). Constraint metrics for genetic variation indicated that *ZNF512B* is intolerant to heterozygous loss-of-function variation (pLI score = 1.000) (**Figure 4D**). Notably, more than half of the RRKQxxP NIM-containing proteins identified from our analysis (**Table 1**) were also intolerant to loss-of-function, indicating their essential role in humans (**Figure 4D**). We then predicted the impact of missense SNPs in ZNF512B using Polyphen, and observed that the ZF SNPs were more functionally damaging (*p*<0.0001, **Figure 4E**), whereas analysis using SIFT showed the ZF residues were also more intolerant to variation (*p*<0.0001, **Figure 4F**).

**Figure 4.**
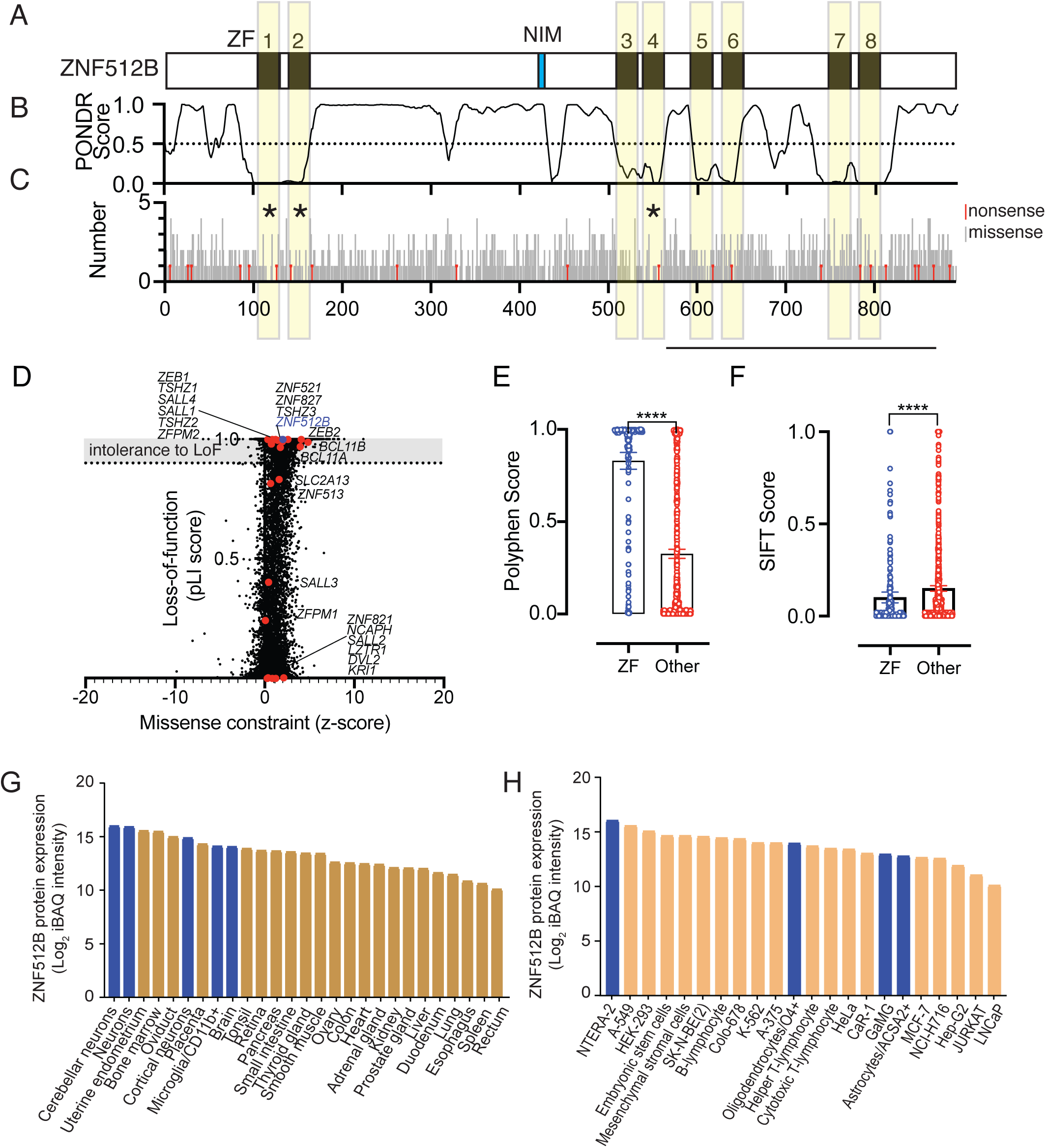
ZNF512B is a neuronal-expressed C2H2-ZF type protein which is intolerant to loss-of-function mutation. **(A)** Schematic depicting the domain organisation of ZNF512B protein; four tandem arrays of atypical and typical C2H2 ZFs are indicated along with an internal NIM. **(B)** Plot of disordered regions as determined by PONDR analysis (0-0.5, ordered; 0.5-1.0, disordered). **(C)** Distribution and frequency of missense (grey) and nonsense (red) SNPs. Asterisks indicate ZFs where less missense SNPs are observed than expected (*p*<0.05, Chi-squared test). **(D)** Constraint metrics of *ZNF512B* nonsense and missense SNPs for all RRKQxxxP NIM-containing proteins (Table 1, red dots) including *ZNF512B* (blue dot). Intolerance to heterozygous loss-of-function mutation is shown with the pLI score (y-axis), with >0.9 – 1.0 indicated as intolerant (grey shading); intolerance to missense mutation is shown (z-score, x-axis). **(E-F)** Analysis of the functional impacts of ZNF512B ZF-specific missense SNPs using Polyphen **(E)** and SIFT analysis **(F)**. Data represents mean±SD, with statistical significance determined using the Mann-Whitney U-test (****, p<0.0001). **(G-H)** Bar graphs showing log2 of iBAQ (intensity-based absolute quantification) values for ZNF512B protein expression across different tissues **(G)**, and cell types **(H)**. Bars highlighted in blue are of brain or neural origin.

**Table 2:** Significant de-enrichment of missense SNPs occurs in specific ZFs of ZNF512B. Missense SNPs occurring in *ZNF512B* were obtained from GnomAD database, and amino acid co-ordinates for each ZNF512B ZF was obtained from UniProt. The expected SNPs were calculated from the 1137 missense SNPs detected in ZNF1512B (892 aa in size). Statistical significance (*p*<0.05, bold) was determined with Chi-squared analysis.

| ZF | Co-ordinates (aa) | Size (aa) | Missense SNPs (Observed) | Missense SNPs (Expected) | $p$-value |
| --- | --- | --- | --- | --- | --- |
| 1 | 105-129 | 25 | 17 | 32 | **0.0303** |
| 2 | 140-163 | 24 | 15 | 31 | **0.0172** |
| 3 | 510-533 | 24 | 21 | 31 | 0.1607 |
| 4 | 540-563 | 24 | 14 | 31 | **0.0105** |
| 5 | 594-618 | 25 | 25 | 32 | 0.3477 |
| 6 | 630-653 | 24 | 33 | 31 | 0.7992 |
| 7 | 750-774 | 25 | 34 | 32 | 0.8027 |
| 8 | 784-807 | 24 | 22 | 31 | 0.2111 |
| All ZFs |  | 195 | 181 | 249 | **0.0003** |

We next compared orthologous ZNF512B protein sequences at varying evolutionary distances demonstrate that ZNF512B is an evolutionarily conserved protein with greater than 80% similarity in mammals compared to humans (**Supplementary Figure 3B**). Importantly, the NIM is conserved in all orthologues indicating that it is functional important domain for organismal function (**Supplementary Figure 3C**). To gain an insight into the expression pattern of ZNF512B protein across different human tissues and cell types, we obtained quantitative mass spectrometry-based proteomics data from ProteomicsDB, an online multi-omics and multi-organism resource^35^. Our analysis revealed that ZNF512B protein is expressed in diverse human tissues and cell types (**Figure 4G & H**). Notably, it is highly expressed in specific cell types within the brain such as cerebellar and cortical neurons and microglia (**Figure 4G**). For cell types, ZNF512B is more abundantly expressed in human pluripotent embryonal carcinoma (NTERA-2) cells, embryonic and mesenchymal stem cells, as well as HEK293 cells (**Figure 4H**).

### ZNF512B-NuRD complex may regulate neuronal-specific gene transcription program

As a ZNF512B is a C2H2-ZF-containing protein interacting with the NuRD complex, we reasoned that it will likely possess a transcriptional regulatory function. To provide better understanding of the role of ZNF512B in gene regulation, we performed RNA-seq analysis on HEK293T cells after shRNA-mediated *ZNF512B* knockdown. To first establish efficient depletion of *ZNF512B*, we designed two shRNA targeting *ZNF512B* in a pLKO.1-based lentiviral vector also encoding an transgene cassette containing an eGFP reporter. The two independent shRNAs targeted either the coding sequence (sh1) or the 3′ UTR (sh2) of *ZNF512B* (**Figure 5A**), with a non-targeting shRNA used as a control (shCTRL). Transduction efficiency for all shRNAs was above 98% (**Figure 5B**). Efficient depletion of *ZNF512B* was confirmed at both the mRNA (**Figure 5C**) and protein level (**Figure 5D**). Principal component analysis of RNA-Seq data showed that *ZNF512B* knockdown replicates were separated in different clusters on both principal components (**Supplementary Figure 4A**). This indicates that gene expression profiles were consistent between replicates but varied between conditions. In total, 1,486 differentially expressed genes (DEGs) were detected in shRNA1-mediated *ZNF512B*, wherein 987 (66%) were downregulated and 499 (34%) were upregulated (*p*-adj <0.05 & fold change >2) (**Figure 5E, Supplementary Table 5**). We observed similar proportions of DEGs in shRNA2-mediated *ZNF512B* knockdown (1,246 total DEGs, with 875 (70%) downregulated and 371 (30%) upregulated) (**Figure 5E, Supplementary Table 5**). To confirm that the gene regulatory changes triggered by each *ZNF512B* shRNA were similar we compared the fold changes of DEGs in sh2 samples then displayed corresponding gene expression changes in sh1 (**Supplementary Figure 4B**). For the majority of the DEGs, fold changes were found similar between sh1 and sh2 suggesting that *ZNF512B* knockdown triggers similar transcriptional changes irrespective of the shRNA used. Overall, 389 DEGs were common to both shRNAs (**Figure 5F)**. Gene ontology analysis of these common DEGs showed significant enrichment for biological processes related to ion transmembrane transport, featuring gene families for voltage-gated calcium channels (CACNA), potassium channels (KCN) and sodium channels (SCN). Notably, Wnt signaling, synaptic function and assembly, and related neuronal processes were also enriched (**Figure 5G**), despite the experiment being performed in HEK293T cells. Importantly, a large proportion of genes involved in these neural-specific processes appear to be significantly downregulated upon *ZNF512B* knockdown (**Figure 5H**), suggesting ZNF512B normally positively regulates many genes involved in neuronal processes.

**Figure 5.**
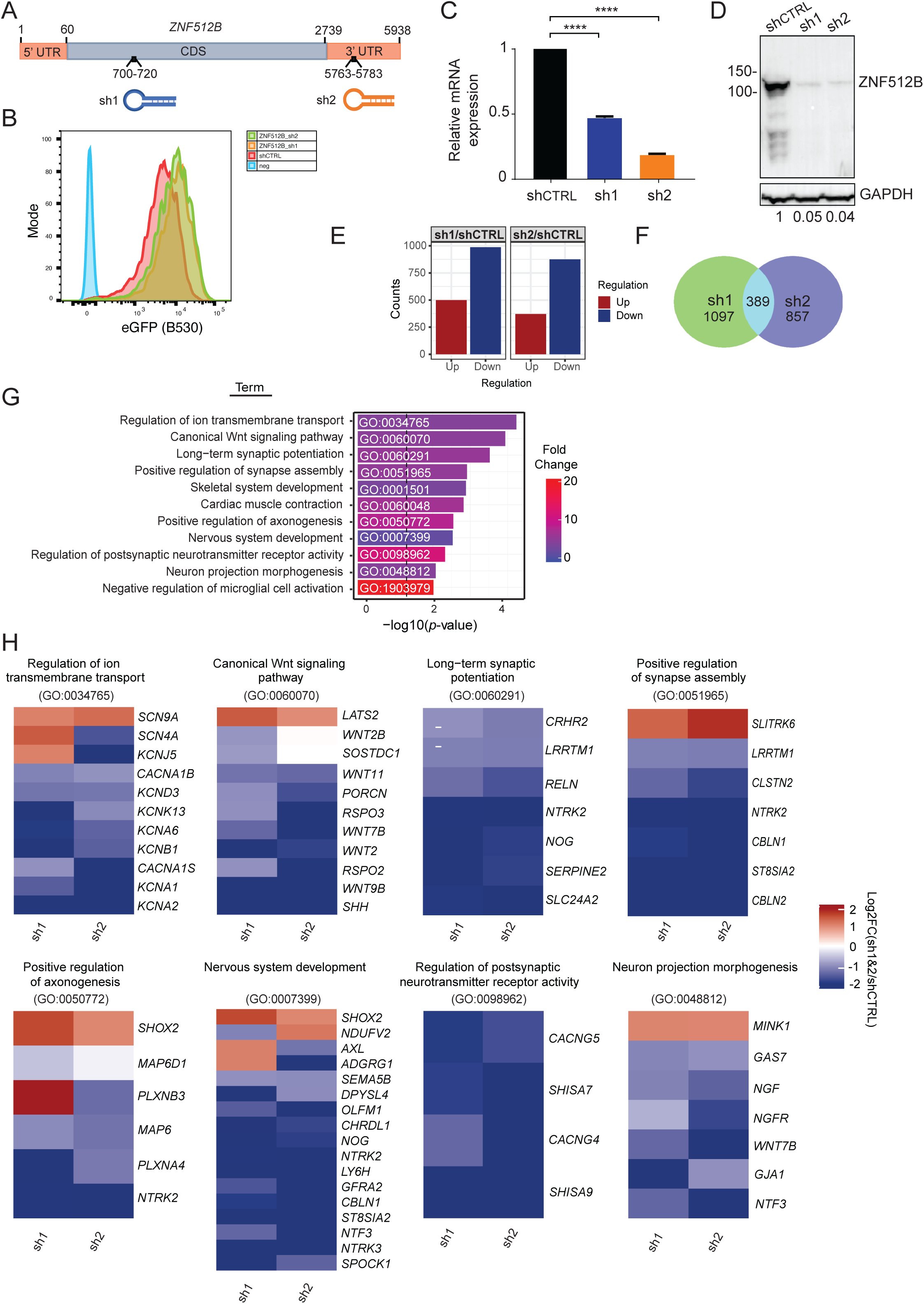
Downregulation of neuronal development genes after ZNF512B depletion. **(A)** Schematic of *ZNF512B* mRNA and location of the shRNA targeting sequences. **(B)** Histogram overlays of eGFP expression in HEK293T cells stably transduced with lentiviral vectors expressing *ZNF512B* shRNAs or a non-targeting shRNA (shCTRL). **(C-D)** Expression of ZNF512B after shRNA knockdown in HEK293T cells, as measured by **(C)** qRT-PCR, with *ZNF512B* gene expression normalized to housekeeping gene *GAPDH*; and **(D)** Western blot for ZNF512B protein expression. GAPDH was used a loading control; values under graph represent densitometric values. **(E**) A barplot showing counts of DEGs in *ZNF512B* shRNA-knockdown-treated cells for two different shRNAs. **(F)** Venn diagram showing number of unique and common DEGs between the two *ZNF512B* shRNAs. **(G)** A bar plot showing enriched gene ontology (GO) terms for biological processes; the dotted line is drawn at *p-*value = 0.05. **(C)** Heatmaps showing expression levels of genes involved in different neuronal processes and pathways identified in the GO analysis.

### ZNF512B is highly expressed in neuronal progenitor cells

Human neural NTERA-2 (herein referred to as NT2) cells were chosen as the model cell line to examine ZNF512B neuronal function as they also can be differentiated into NPCs. Firstly, to determine if ZNF512B is an essential factor for cell proliferation and survival in NT2 cells, we depleted *ZNF512B* expression using shRNA knockdown. In NT2 cells, transduction efficiency for all shRNAs was above 90% (**Figure 6A**). NT2 cell proliferation was then analysed over 5 days via MTT (3-(4,5-dimethylthiazol-2-yl)-2,5-diphenyltetrazolium bromide) assays. The assays showed that NT2 cell proliferation was inhibited as early as 24 h post-*ZNF512B* depletion, while NT2 cells transduced with shCTRL exhibited robust proliferation (**Figure 6B**). We further examined the significance of ZNF512B in cell survival using clonogenicity assays because NT2 cells possess cancer stem cell-like features. ZNF512B-depleted cells significantly lost their colony-forming ability compared to shCTRL (**Figure 6C**). Taken together, these results, confirm that ZNF512B is an essential factor in NT2 cell proliferation and survival.

**Figure 6.**
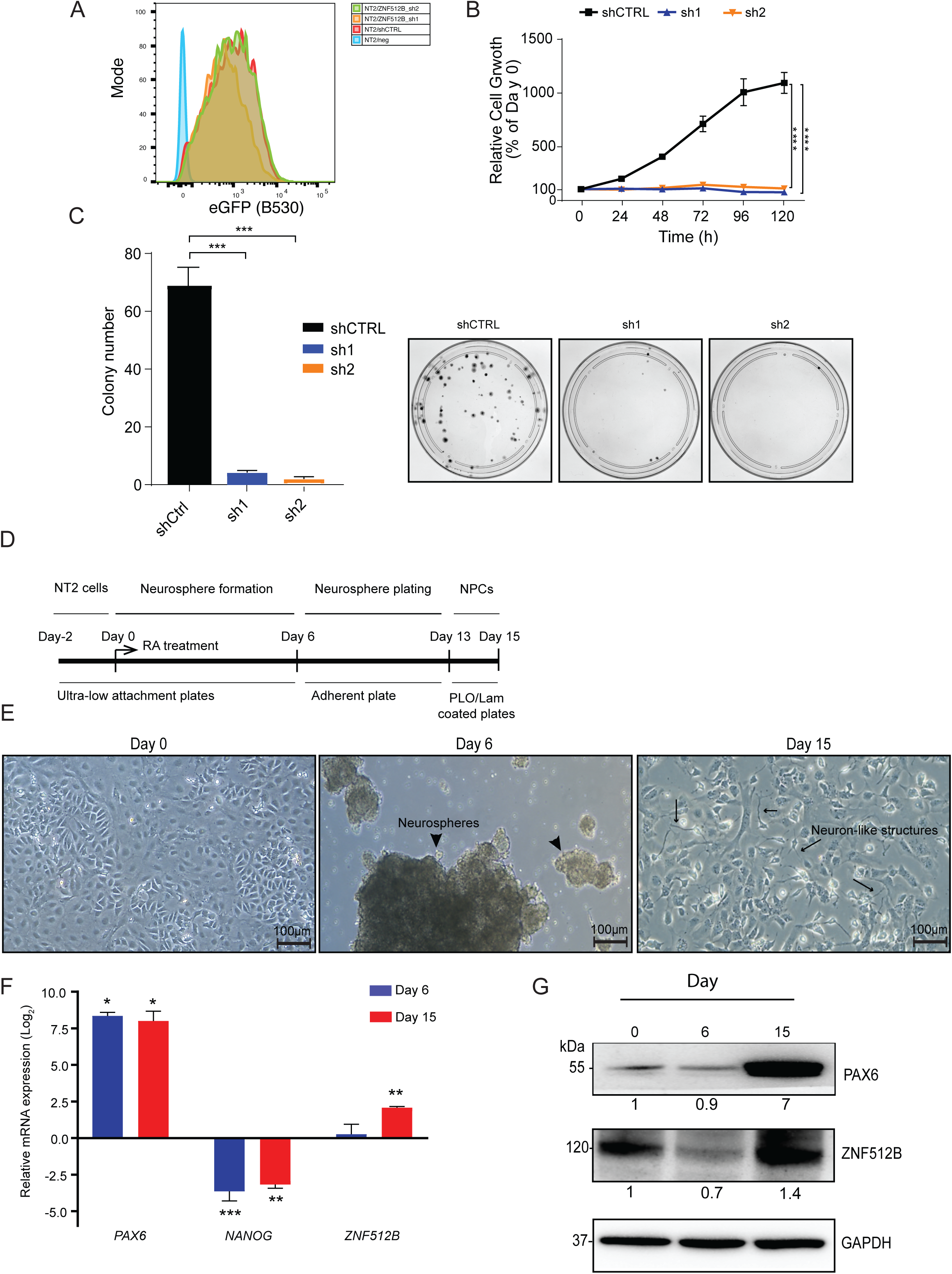
ZNF512B expression is important for neural cell growth and differentiation. **(A)** Histogram overlays of eGFP expression in NT2 cells stably transduced with lentiviral vectors expressing *ZNF512B* shRNAs or a non-targeting shRNA (shCTRL). **(B-C)** Measurement of **(B)** NT2 cell proliferation (line graph), and **(C)** colony forming ability (bar graph, *left panel*) after *ZNF512B* shRNA knockdown. Representative images of colonies in plates after *ZNF512B* knockdown **(C**, *right panel*). Data is represented as the mean ± s.e.m for 3 experiments each performed in triplicate. Statistical analysis was performed using a two-way ANOVA and unpaired t-test for proliferation and clonogenicity assays, respectively (***p<0.001; ****p < 0.0001). **(D)** Schematic diagram showing the timecourse of the NT2 neural differentiation assay performed over 15 days (RA: retinoic acid; NPC: neural progenitor cell; PLO/Lam: poly-L-ornithine/laminin). **(E)** Representative micrographs of NPC differentiation; scale bars =100 μm captured at three different time points; arrowhead=neurospheres; arrows=neuron-like structures. **(F)** Bar graph illustrating qRT-PCR data. Expression of *NANOG*, *PAX6*, and *ZNF512B* were profiled at key timepoints. Gene expression was normalised to Day 0. Data represents mean ± s.e.m of 3 independent experiments performed in triplicate. Statistical analysis was performed using one-way ANOVA (*p< 0.05, **p < 0.01; ***p < 0.001). **(G)** A representative Western blot showing increased protein levels of PAX6 and ZNF512B in NPCs.

Our transcriptome data demonstrated that the ZNF512B-NuRD complex could play an essential role in neurogenesis. Thus, we hypothesised that ZNF512B might be involved in gene regulation during NPC differentiation. NPCs are a crucial population of early neuronal cells which can give rise to many neuronal cell types (e.g., dopaminergic neurons and oligodendrocytes) that constitute the central nervous system in brain^36^. To provide evidence for our hypothesis, we examined the expression pattern of ZNF512B in undifferentiated NT2 cells and NPCs. To test this, we differentiated NT2 cells into NPCs in the presence of retinoic acid and collected cells at different timepoints, namely, 0, 6, and 15 days (**Figure 6D**). Brightfield images demonstrate successful differentiation of stem-like NT2 cells (day 0), into neurospheres (day 6) and NPCs (day 15) (**Figure 6E**). We then performed qRT-PCR analysis on total RNA extracted at different days (**Figure 6F**). Efficient differentiation of NT2 cells into NPCs was demonstrated using *NANOG* and *PAX6* as stem cell and NPC marker genes, respectively. As expected, qRT-PCR analysis indicated a marked decrease in *NANOG* and increase in *PAX6* expression during NPC formation (**Figure 6F**). Surprisingly, the expression of ZNF512B at both mRNA and protein levels was upregulated in NPCs when compared to days 0 and 6, resembling a NPC-specific gene expression pattern (**Figure 6G**). This observation further reinforces that ZNF512B might play a crucial role in the positive regulation of neuronal gene expression programs during differentiation into NPCs.

## Discussion

Several genome-wide association studies have identified *ZNF512B* as a disease-susceptibility gene for ALS^26,27^. The presence of a non-coding single nucleotide variant in *ZNF512B* (rs2275294) has been shown to correlate with lower mean survival^28,29^. The rs2275294 variant, residing in a putative enhancer region of *ZNF512B*, downregulated *ZNF512B* transcription, and as a consequence, influenced TGFβ pathway signalling, a pathway critical for protecting brain motor neurons against damage and degeneration^26,37^. These observations provided preliminary evidence supporting a potential role of ZNF512B in neurogenesis and neuroprotection.

Here we show that ZNF512B depletion leads to a marked dysregulation of neural-related molecular and cellular pathways. On a mechanistic level, we suggest that ZNF512B regulates neural-specific transcription programs through interaction with the RBBP4-NuRD complex. RBBP4 is found in several multi-subunit protein complexes involved in chromatin remodelling and gene transcription. These complexes include NuRF^38^, PRC2^39^, CAF-1^40^, Sin3^41^, and HAT1^42^. Exclusive interaction of ZNF512B with the RBBP4-NuRD complex is explained by the presence of the NIM in ZNF512B, which has been shown to mediate binding of auxiliary factors to the RBBP4-NuRD complex only^43,44^. This observation is consistent with the previous large-scale protein-protein interaction studies, where enrichment of ZNF512B was observed in NuRD subunit pulldowns^15,17^.

There is a growing body of evidence describing differential roles of the NuRD complex in neurodevelopment^45^. For example, in mouse knockout studies, MBD3 depletion significantly reduced cortical thickness and disrupted transcriptional circuits, with effects propagated on neuronal cell-lineage decisions and brain developmental transitions^46,47^. Similarly, CHD4/NuRD activity regulates a network of genes required for appropriate synaptic assembly and transmission, greatly influencing neuronal connectivity in the mammalian cerebellar cortex^19,48^. The importance of the NuRD complex in neurodevelopment is also highlighted in mutational studies where pathogenic variants in genes encoding NuRD subunits *CHD3/4* and *GATAD2B* have led to macrocephaly and intellectual disabilities^49–52^. Our data indicate that the assembly of ZNF512B into the NuRD complex may constitute the specialised transcriptional machinery that defines a neural-specific transcriptional program. However, our findings do not distinguish between NuRD-dependent and -independent functions of ZNF512B, making it a limitation of this study. Depending on the availability of high-quality antibody reagents, genome-wide occupancy studies using ChIP-Seq for core DNA-binding subunits of the NuRD complex (e.g., MBD3), along with ZNF512B, in the promoter region of neural genes would help define NuRD dependency based on occupancy. Interestingly, a recent study demonstrated a role for ZNF512B in binding ‘TTC’ DNA sequences which are arranged in a repetitive or non-consecutive manner in pericentric repeats regions^53^. This is an important early step in heterochromatin formation. Furthermore, CHD3 and CHD4 can form distinct NuRD complexes enriched in perinuclear heterochromatin^54^. NuRD-independent roles for ZNF512B in transcriptional regulation have not been properly defined, with the sole evidence from ZNF512B overexpression studies with GFP-tagged normal or K423 mutant (non-NuRD-binding) ZNF512B^30^.

NTERA-2 human pluripotent embryonic carcinoma cells are frequently used as an *in vitro* cell model to produce both progenitor and mature neuron-like cells^55,56^. NPCs are an important population of cells in the central nervous system as they give rise to distinct cell types (e.g., glia, and dopaminergic neurons)^36^. We observed that *ZNF512B* depletion prevented normal NT2 cell growth and proliferation, which would make it impossible to differentiate NT2 cells into NPCs in the absence of ZNF512B. Therefore, we studied ZNF512B expression pattern at both mRNA and protein levels during NPC formation. Our data showed a marked upregulation of ZNF512B expression in NPCs, matching the pattern of PAX6 expression. Notably, in our study, NPCs were not enriched and may still contain undifferentiated or partially differentiated cells. Thus, we speculate that the increase in ZNF512B expression would have been more pronounced if NPCs were specifically enriched. Upregulation of ZNF512B in NPCs, supports an essential role for ZNF512B in NPC formation, but also later for specific neuron-subtype formation. Future work should explore ZNF512B expression in mature neurons such as dopaminergic neurons because they are widely studied in ALS and Parkinson’s disease models.

H2A.Z, a variant of H2A, is a key determinant of NPC differentiation through regulating the expression of neural genes^57,58^. It has been shown that the NuRD complex facilitates replacement of H2A with H2A.Z in the promoter region of activity-dependent genes in the mouse brain^20^. Previous H2A and H2A.Z containing mononucleosome-specific protein-protein interaction studies have reported the association of ZNF512B with H2A.Z-containing mononucleosomes, but not H2A-containing mononucleosomes^59,60^. This suggests that the ZNF512B-NuRD complex may specifically recognize H2A.Z-containing nucleosomes positioned in the promoter region of neural genes. In this model, ZNF512B would recruit the NuRD complex to H2A.Z-containing nucleosomes and define a specific gene expression program during NPC formation. These observations will pave the way to study the dynamic of gene expression program during neuronal cells differentiation with a focus on the ZNF512B-NuRD-H2A.Z axis.

## Materials and Methods

### Plasmid constructs

The wildtype *ZNF512B* coding sequence (obtained as a cDNA from Horizon Discovery: MHS6278-211690838) was cloned into a HA epitope-containing pcDNA3.1(+) vector using a Gibson assembly approach. NIM (RRKQxxP) motif deletion and alanine substitutions were introduced using Q5 (NEB)- and QuikChange (Agilent)-mediated site-directed mutagenesis, respectively. A list of primers used for cloning are available upon request. FLAG-tagged constructs expressing NuRD subunits (CHD4-C, MBD3, MTA2, RBBP4 and GATAD2B) were a kind gift from Professor Joel Mackay, The University of Sydney^14^. For gene knockdown, two different short hairpin RNA oligos **(Supplementary Table 4)** targeting *ZNF512B* mRNA containing *AgeI/EcoRI* compatible ends were cloned into pLKO.1-based vectors co-expressing green fluorescent protein (GFP) linked via picornaviral 2A peptide sequence to puromycin. *Arabidopsis thaliana miR159*a cloned into the same plasmid backbone was used as a non-targeting control (shCTRL).

### Cell culture, transfection, and transduction

All human cell lines used in this study have been authenticated by short tandem repeat profiling (Cellbank, Australia). Suspension-adapted human embryonic kidney cells (Expi293F™) were grown in Expi293F Expression Medium (Thermo Fisher Scientific, Waltham, MA, USA) on a horizontal orbital shaker (130 rpm) placed inside a humidified incubator at 37 °C, 5% CO2. For Co-IP experiments, Expi293F cells were grown to a density of 1.5-2×10^6^ cells/mL and plasmid DNA (2 μg) was transfected into the cells using linear polyethylenimine (PEI; 8 μL, 1 mg/mL). After 72 h, cells were harvested, washed twice with cold PBS, centrifuged (300 *g*, 5 min), snap-frozen in liquid nitrogen and stored at −80 °C until further use. NTERA-2 (also known as NT2) cells and HEK293T cells were grown in DMEM/F12 (Thermo Fisher Scientific) supplemented with 10% FCS (v/v), penicillin (100 U/mL) and streptomycin (100 μg/mL). For transductions, lentivirus was produced by calcium phosphate transfection of HEK293T cells with packaging plasmids (pRSV-Rev, pMDLg/p.rre and pMD2.VSV-G) and *ZNF512B* shRNA knockdown vectors. Viral supernatant was collected after 48 h, 0.45 μM-filtered, snap-frozen in liquid nitrogen and stored at −80 °C. NT2 cells were grown to a confluency of 0.35×10^6^ cells/mL in a 6-well plate before addition of fresh media containing viral supernatant and Polybrene (8 μg/mL; Sigma). The plate was centrifuged at 1,500 rpm for 1.5 h and then incubated at 37 °C for 48-72 h. Cells were collected and checked for GFP expression using flow cytometry.

### Co-immunoprecipitation

Equimolar amounts of FLAG- and HA-tagged pcDNA3.1 plasmids were co-transfected into Expi293F cells using linear polyethylenimine (PEI; 8 μL, 1 mg/mL). After 72 h, cells were harvested, washed with PBS and resuspended in lysis buffer (600 μL; 50 mM Tris-HCl, 250 mM NaCl, 1x EDTA-free SIGMAFAST protease inhibitor cocktail (PIC) (Sigma), 1 mM PMSF, 0.2 mM DTT, pH 7.5). Input (10% of total volume) was collected, and the remaining lysate was mixed with 25 μL of pre-washed anti-HA Sepharose 4B beads (Sigma-Aldrich) on a rotator (25 rpm) for 2 h at 4 °C. Following incubation, beads were washed with 5 times with cold wash buffer (1 mL; 50 mM Tris-HCl, 200 or 500 mM NaCl, 0.5% (v/v) IGEPAL® CA630, pH 7.5). Finally, three rounds of elution were performed by adding 30 μL of a 150 μg/μL stock of 3xFLAG peptide (#F4799, Sigma-Aldrich) and incubating samples in a thermomixer (800 rpm) for 40 min at 4°C. Western blot analysis was carried out as previously described^61^. Antibodies used here are as follows: anti-FLAG (1:5,000 dilution; #A8592-1, Sigma-Aldrich), anti-HA (1:5,000 dilution; #2999, Cell Signaling Technology).

### Affinity purification followed by LC-MS/MS analysis

For protein-protein interaction studies, label-free quantitative mass spectrometry experiments were conducted on affinity-purified HA-ZNF512B samples (n=3). In brief, Expi293F cells were transfected with either pcDNA3.1-HA-ZNF512B or -HA-ZNF512B mutants or control pcDNA3.1-HA-only (empty vector) plasmids. After 72 h, collected cells were subjected to fractionation (1 mL; 10 mM HEPES-KOH, 1.5 mM MgCl_2_, 10 mM KCl, 5% Glycerol (v/v), 1x PIC, 1 mM PMSF, 1 mM DTT, pH=7.9). Nuclear pellets were resuspended in 500 μL lysis buffer (50 mM Tris-HCl, 150 mM NaCl, 1% (v/v) Triton X-100, 1x protease inhibitor cocktail, 1 mM PMSF, 1 mM DTT, 25-50 units Benzonase, pH 7.5), sonicated and centrifuged (16,000 g, 10 min) before mixing with pre-washed anti-HA beads (Sigma-Aldrich) on a rotator (25 rpm) for 2 h at 4°C. Bound proteins were washed twice with 1 mL cold wash buffer (50 mM Tris-HCl, 200 mM NaCl; 0.5% IGEPAL) and twice with 1 mL TBS (50 mM Tris-HCl, 200 mM NaCl). On-bead tryptic digestion, peptide desalting, and mass spectrometry sample loading were performed as described previously^14^.

The Q-Exactive HFX mass spectrometer was set to a data-dependent acquisition mode (DDA). In the DDA run, each full scan MS1 was operated at a resolution of 70,000 at 200 m/z and 100 ms injection time. The top 15 most intense precursor ions were selected to be fragmented in the Orbitrap via high-energy collision dissociation activation. Raw data files were analysed in MaxQuant (version 1.6.6.0) using standard settings, where methionine oxidation (M), and carbamidomethyl cysteine (C) were selected as variable and fixed modifications, respectively. Statistical analysis of the generated text file was carried out in R Studio using the “DEP” package. Significance was drawn at Log_2_Fold change >1 and *p*-value <0.05. Visualisation of results was performed using the R package “Enhanced volcano” (github.com/kevinblighe/EnhancedVolcano).

### Motif and SNP analysis

NIM motifs RRKxxxP, RKxxxPxK, RRKQxxP were used as the search queries to retrieve human proteins containing those motifs using Scansite 4.0. Lists of proteins were generated (**Supplementary Table 1**) and these were manually curated for NuRD complex subunit interactions and subcellular localization (BioGRID), for direct involvement or association with genetic disease (OMIM), and for essentiality revealed by CRISPR/Cas9 screen dependency (Human DepMap, version 24Q3). Using the DAVID bioinformatics resource, ontology and protein-protein interaction enrichment was performed. Sequence logos generated from multiple sequence alignments of enriched motifs were generated using WebLogo. To visualize the distribution of specific NIM across the length of proteins, we computed kernel density estimates of motif positions and projected them on a relative scale from 0% to 100% of the protein sequence. A custom R function was used to calculate and interpolate density values at specified intervals along the protein length. Briefly, motif start positions were extracted from protein sequences and passed to the density() function in R with appropriate smoothing parameters (e.g., bandwidth). The resulting kernel density estimates were interpolated at regular intervals (e.g., every 1%) using approx() to obtain smoothed density values along a normalized x-axis representing the relative position within the protein. These values were visualized using a heatmap, with color intensity corresponding to motif density. The scale bar reflects density values, enabling comparison of motif enrichment patterns across protein regions.. Gene ontology analysis for NIM-containing motifs was performed using DAVID. *ZNF512B* SNPs primarily focusing on missense and nonsense variants were retrieved from gnomAD, as well as constraint metrics signifying intolerance to loss of function (pLI score) and missense constraint (z-score). The functional impact of missense mutations were analysed using Polyphen-2 and SIFT algorithms. Prediction of protein disorder was performed using PONDR with a score of 0 to 0.5 indicating an amino acid residue was located in an ordered region, and 0.5 to 1.0 indicating a disordered region.

### Structure Prediction and Modeling

To evaluate the potential interaction between ZNF512B and individual NuRD complex subunits, we performed pairwise structural modeling using AlphaFold-Multimer v2.3 via the ColabFold platform (https://colab.research.google.com/github/sokrypton/ColabFold). This tool integrates multimeric modeling capabilities with deep learning-based structural inference. Confidence in predicted interactions is assessed using the interface-predicted template modeling (ipTM) score, where values above 0.8 indicate high-confidence predictions and scores below 0.6 suggest unreliable prediction^62^. To further validate the structural robustness of the model, Predicted Aligned Error (PAE) and predicted Local Distance Difference Test (pLDDT) scores were analysed. Low inter-chain PAE values support reliable interface geometry and high pLDDT values indicate strong confidence in the modelled structures.

### 3D modeling and binding analysis

The 3D model of the ZNF512B-RBBP4 interaction complex was constructed using the experimentally validated complex of RBBP4 bound to BCL11A peptide^63^ available in the Protein Data Bank (PDB: 5VTB). The ZNF512B NIM motif plus flanking residues (420-434) has the amino acid sequence “HRRKQKTPKKFTGEQ” whereas the BCL11A peptide has 15 amino acid residues (2-16) with the sequence “SRRKQGKPQHLSKRE”. To this end, the latter was mutated to match the ZNF512B peptide with Histidine inserted at the starting position using the ‘Build and Edit Protein’ protocol in the Biovia Discovery Studio software suit (DS2022). Following this, the ZNF512B peptide and the RBBP4 protein were subjected to energy minimization to attain their native state using CHARMm36 force fields in DS 2022. The ‘Minimization’ protocol with the steepest descent algorithm for a maximum of 2000 steps without an implicit solvent model and a non-bonded lower cut-off distance of 10 Å was used.

By mutating each amino acid residue of ZNF512B peptide into alanine iteratively, 15 mutated models were prepared with ‘Build and Edit Models’ protocol in DS 2022. The impact of the mutation on the binding affinity of ZNF512B with RBBP4 was calculated using ‘The Calculate Mutation Energy (Binding) protocol’ (available in DS 2022) where the energy effect of each mutation on the binding affinity (mutation energy, ΔΔGmut) is measured as the difference between the binding free energy in the mutated structure and wild-type protein: ΔΔGmut=ΔΔGbind (mutant)-ΔΔGbind (wild type). The binding free energy, ΔΔGbind, is the difference between the free energy of the complex and unbound state of the interacting components. All energy terms are calculated using CHARMm36 force field, and the electrostatic energy is calculated using a Generalized Born implicit solvent model. The total free energy is calculated as an empirical weighted sum of van der Waals (EvdW) interaction, electrostatic interactions (ΔGelec) including the effect of ionic strength I in a pH-dependent mode, an entropy contribution (-TSsc) related to the changes in side-chain mobility, and a non-polar, surface-dependent contribution to solvation energy (ΔGnp) for each of the components in the complex using the following equation: ΔGtot(pH) = aEvdW+ ΔGelec(pH,I)-cTSsc+ ΔGnpwhere a and c are empirical scaling parameters with the optimal value of a≈0.5 and c≈0.8.

### Molecular Dynamics Simulation

Molecular dynamics simulations were conducted in a vacuum environment to explore the stability and folding, conformational changes, and dynamics in forming various electrostatic interactions using the ‘Standard Dynamics Cascade’ protocol of DS2022. The CHARMm36 force-field was applied to all systems with initial energy minimization of 1000 steps with the steepest descent and 2000 steps with Adopted Basis NR with the final RMS gradient of 0.1. After minimization, all the systems were subjected to a heating step where the initial temperature was raised from 50 K to 300 K in 50 ps time intervals, followed by an equilibration step for another 50 ps. Finally, the production run was carried out for 20 ns in an NVT assembly (normal volume and temperature) at a constant temperature of 300 K, and the results were saved after every 0.02 ns. All the trajectories were analysed for parameters like root-mean-square deviation (RMSD) and bond interaction between the wild type and all the mutated models of the ZNF512B-RBBP4 complex using the ‘Analyse Trajectory Protocol’ of DS 2022.

### NT2 cell culture and neural differentiation

NT2 cells were seeded at a density of 1×10^6^ cells in ultra-low attachment plates and incubated in 5 mL of DMEM/F12 media containing 2-mercaptoethanol (50 μM) for 48 h at 37 °C. Retinoic acid (10 μM) was then added to induce differentiation (day 0) and cells were left undisturbed for 6 days. On day 7, formed clusters were transferred to adherent T75 flasks and subsequently cultured for another 6 days. On day 13, cells were trypsinised and replated on laminin (1 μg/mL; Sigma) and poly-L-ornithine (0.001% (w/v); Sigma)-coated tissue culture dishes for 48 h at 37 °C. Western blots were performed to check protein expression of ZNF512B (1:5,000 dilution; #A303-234A, ThermoFisher) and PAX6 (1:5,000 dilution; #ab5790, Abcam).

### RNA isolation and qRT-PCR analysis

RNA was extracted from cells collected on day 0, 6 and 15 using the TRIzol reagent (Invitrogen) according to the manufacturer’s instructions. and reverse-transcribed into cDNA using iScript™ gDNA Clear cDNA Synthesis Kit (Bio-Rad). For qRT-PCR reactions, diluted cDNA was mixed with SYBR-green reagent (Bioline) and loaded into a 96-well plate containing primers in distilled water. Ct values were first normalized to GAPDH expression levels, and the relative mRNA expression was calculated using the 2^−ΔΔCt^ method. A list of primers used for qRT-PCR is provided in **Supplementary Table 4**.

### Cell proliferation assays

Cell proliferation was measured by 3-(4,5-methylthiazol-2-yl)-2,5-diphenyltetrazolium bromide (MTT) assay (Merck Millipore). NT2 cells, transduced with either *ZNF512B* shRNA- or shCTRL-containing lentivirus for 72 h, were then plated (3,500/well) in triplicate in a 96-well plate. MTT reagent was added daily at 37 °C overnight over the course of 5 days. A solution mixture of isopropanol/HCl was used to quench the reaction and then absorbance was measured at 570 and 630 nm using a Wallac 1420 Victor plate reader (Perkin Elmer). To assess cell clonogenicity, NT2 cells transduced with either *ZNF512B* shRNA or shCTRL lentiviruses for 72 h (1,500/10 cm dish), were incubated at 37 °C for 14 days. Cells were fixed in ice-cold methanol and stained with Giemsa solution (Sigma-Aldrich).

### RNA-Seq and bioinformatics analysis

For RNA-Seq analysis, HEK293T cells (1×10^6^) were transduced with shZNF512B and shCTRL lentiviruses for 72 h. Total RNA isolation was performed using TRIzol reagent (Invitrogen) according to manufacturer’s instructions. RNA quantitation, integrity check and sequencing were carried out as described previously^64^. For RNA sequencing results, paired-end fastq reads were aligned against the human reference genome (GRCh38 – release101) using STAR aligner (version 2.7.10a)^65^. To perform the differential gene expression (DGE) analysis between shCTRL and *ZNF512B* shRNA samples, first, read counts for each gene were obtained using R function FeatureCount from the R package Rsubread^66^. Genes with sufficient read counts (greater than or equal to 10 in all replicates) were then subjected to DGE analysis using DESeq2^67^. Genes with *p*-adj < 0.05 and log2fold change greater than or equal to 1 were considered differentially expressed. RNA-Seq data visualization was performed using our in-house developed R package (https://github.com/cparsania/parcutils). Gene ontology analysis of differentially expressed genes against all human genes (background) was performed using R package clusterProfiler^68^.

## Acknowledgments

This work was supported by National Health and Medical Research Council Grants (#2037697 to C.G.B. #1128748 & 1177305 to JEJR, #1196405 to U.S.) and by a Cancer Council NSW Project Grant (RG20-12 to U.S. & C.G.B.). This research was supported by Cure the Future for the provision of equipment and scientific services. H.F. was supported with an Australian government Research Training Program PhD scholarship and a Tour de Cure PhD support grant. We acknowledge the technical support we received from Sydney Cytometry and Sydney Mass Spectrometry core facilities at The University of Sydney.

## Data accessibility

All raw MS data have been deposited to the ProteomeXchange Consortium via the PRIDE partner repository with the dataset identifier PXD039013. RNA-seq data will be submitted to GEO upon publication. PDB files for ZNF512B-RBBP4 conformations will be submitted to ModelArchive upon publication.

