## Supplementary Figures for "The ZNF512B-NuRD complex co-ordinates a neural-specific gene expression program"

Supplementary Figure 1

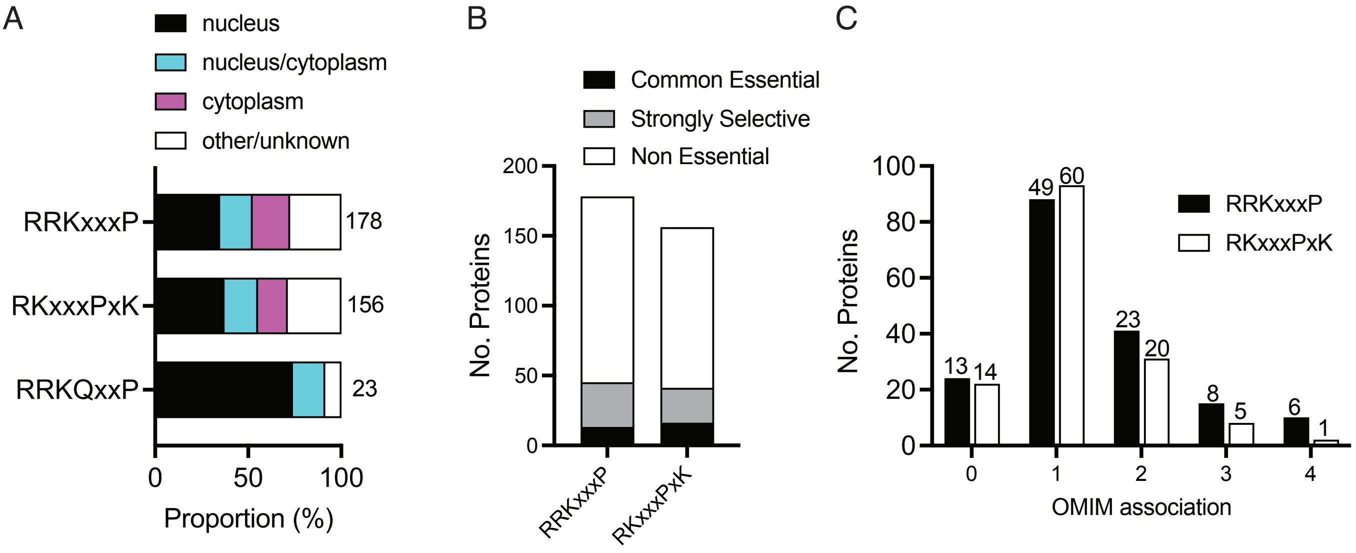

Supplementary Figure 2

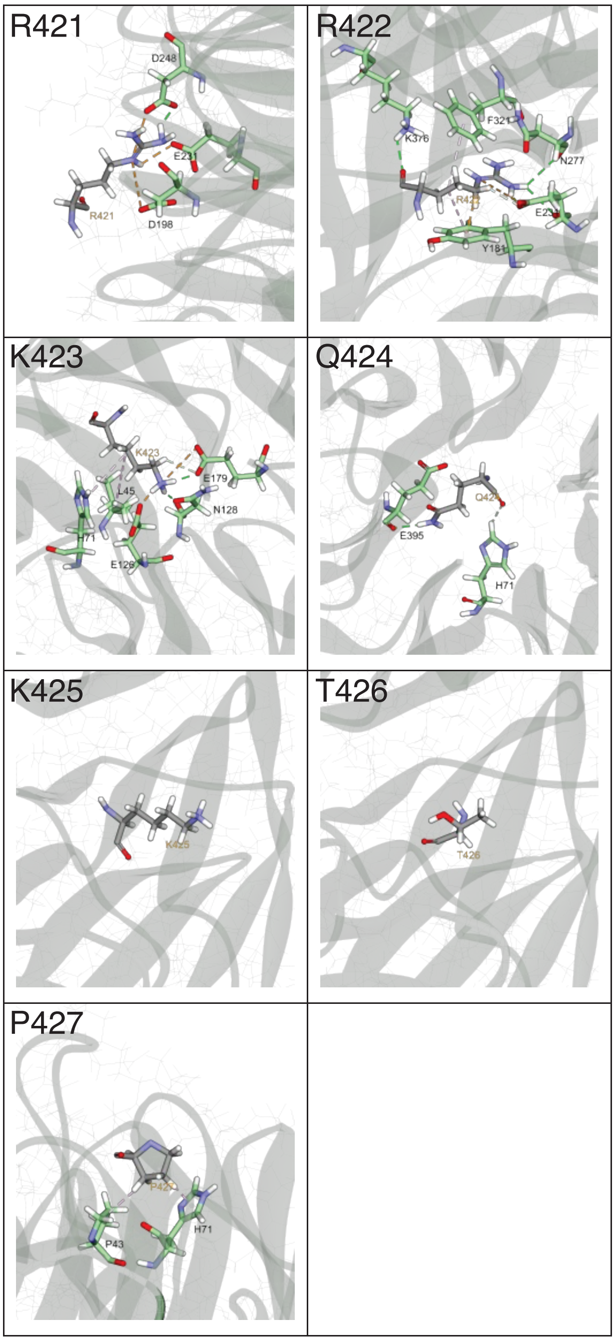


!

Supplementary Figure 3

A

| | | | |
|---|---|---|---|
| **ZF1** | 105 | VKCPNSGCWLEFP-SIYGLKYHYQR-C | 129 |
| **ZF2** | 140 | FPCPF--CEAAFT-SKTQLEKHRIWNH | 163 |
| **ZF3** | 510 | AVCPT--CNVVTRKTLVGLKKHMEV-C | 533 |
| **ZF4** | 540 | LKCQH--CRKQFK-SKAGLNYHTMAEH | 563 |
| **ZF5** | 594 | LRCPQEGCGAAFS-SLMGYQYHQRR-C | 618 |
| **ZF6** | 630 | FPCTH--CGKTYR-SKAGHDYHVRSEH | 653 |
| **ZF7** | 750 | VNCPNDCCEAIYS-SVSGLKAHLAS-C | 774 |
| **ZF8** | 784 | YRCLL--CPKEFS-SESGVKYHILKTH | 807 |

B

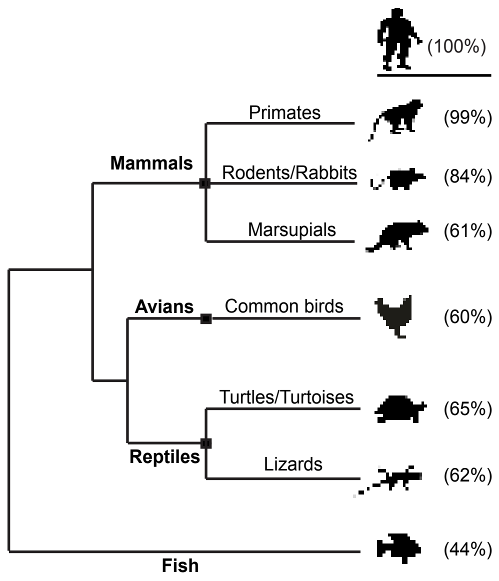


C

NIM

| | | |
|---|---|---|
| **human** | *416* | ERTKHRRKQKTPKKFTGE |
| **chimp** | *416* | ERTKHRRKQKTPKKFTGE |
| **mouse** | *403* | ERTKHRRKQKTPKKFTGE |
| **opossum** | *446* | ERSKHRRKQKTPKKFTGE |
| **chicken** | *325* | ERMKHRRKQKTPKKFTGE |
| **turtle** | *239* | ERMKHRRKQKTPKKFTGE |
| **lizard** | *256* | ERMKHRRKQKTPKKFTGE |
| **zebrafish** | *196* | ERMKHRRKQKTPKKFTGE |

Supplementary Figure 4

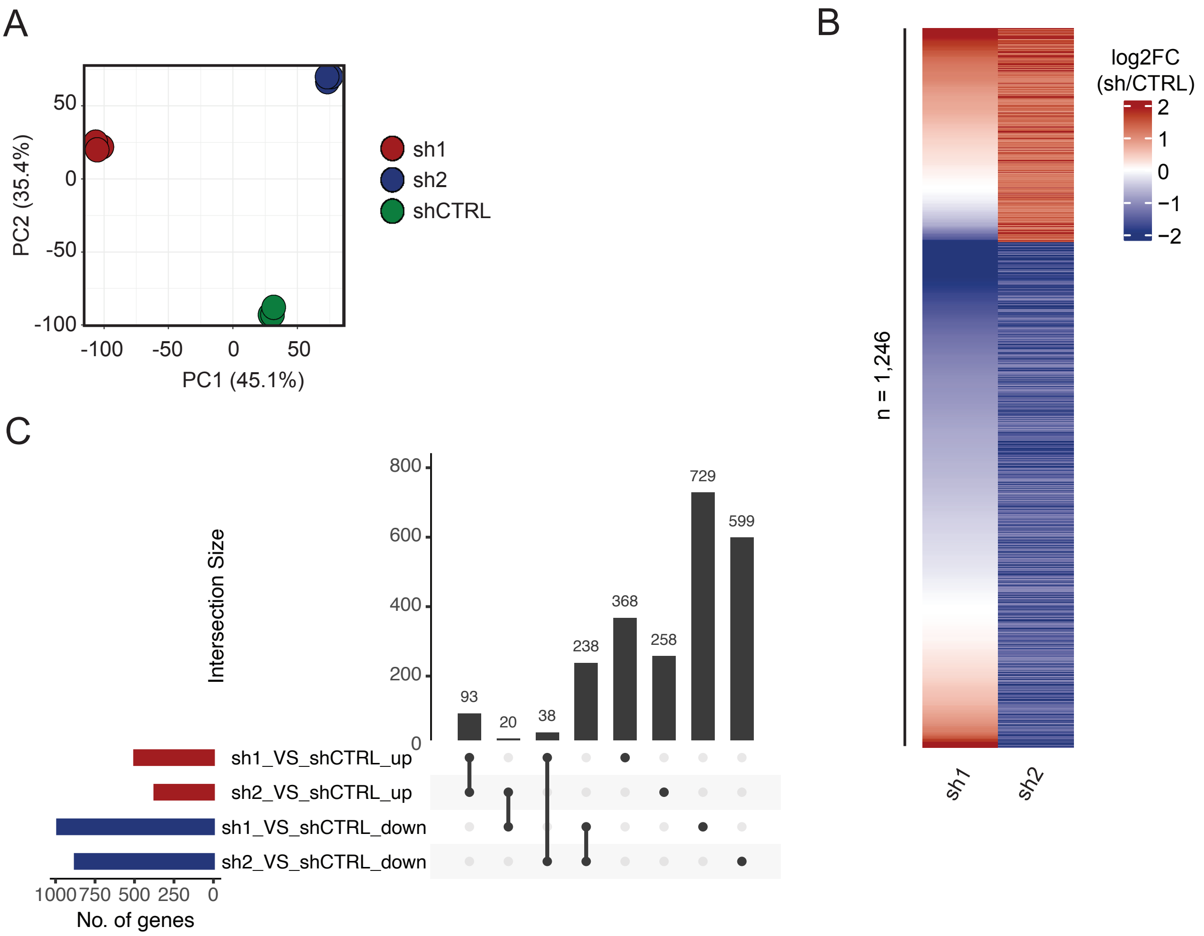
