## Supplementary material for "The ZNF512B-NuRD complex co-ordinates a neural-specific gene expression program": Supplemenary Figure Legends

**Supplementary Figure 1. Cellular and genetic characteristics of NIM-containing proteins, related to Figure 1. (A)** Bar graph showing proportion of NIM proteins with defined subcellular localisation; data from manual curation of NIM proteins from BioGRID. Organelles other than nucleus and cytoplasm including endoplasmic reticulum, Golgi, mitochondria and plasma membrane are listed as 'Other'. **(B)** Cell-line dependency categorisation from the Human DepMap, manually curated for each protein. **(C)** Association of NIM-containing proteins with genetic disease; data from manual curation of NIM proteins.

**Supplementary Figure 2. Key amino acid residues involved in the ZNF512B-RBBP4 interaction, related to Figure 2.** Close-up single amino acid views of residues 421 to 427 which comprise the NIM RRKQxxP of ZNF512B and the molecular interactions made with RBBP4. The type of bonds formed and the chemical groups involved are shown.

**Supplementary Figure 3. Sequence conservation of ZNF512B in multiple species, related to Figure 4. (A)** Amino acid alignment of typical and atypical C2H2 ZFs in ZNF512B; red shading represents Cys residues that co-ordinate C2H2 ZFs; yellow shading represents His residues, orange shading represents the co-ordinating His or Cys residues that distinguish C2H2 and atypical C2H2 (C2HC) ZFs respectively. Each tandem array features one C2HC and C2H2 ZF cluster. **(B)** Amino acid alignment of the NIM motif (blue shading) in ZNF512B in vertebrate species. **(C)** Phylogenetic tree showing ZNF512B protein conservation in vertebrate species. Data on orthologues was acquired from the Ensembl genome database. Values represent the percent identity to the human ZNF512B protein sequence.

**Supplementary Figure 4. Additional analysis of differentially expressed genes in HEK293T cells after *ZNF512B* shRNA knockdown, related to Figure 5. (A)** Principal component analysis of RNA-seq data obtained from HEK293T cells transduced with two independent ZNF512B shRNA or a non-targeting control shRNA (shCTRL). **(B)** Direct comparison of DEGs detected in *ZNF512B* shRNA and level of fold change. All DEGs from ZNF512B sh#2-treated cells were clustered then the fold change of those DEGs in sh#1 were displayed alongside. **(C)** Upset plot displays the degree of unique and shared DEGs detected between *ZNF512B* sh1 and sh2 datasets.

**Supplementary Table 1**. List of human proteins with NuRD interaction motifs (NIMs) obtained from a Scansite 4.0 database search. RRKxxxP, RKxxxPxK and RRKQxxP were used as the search strings, where ‘x’ represents any amino acid. Manual curation was used to obtain adjacent sequences, protein size, NIM amino acid co-ordinates (start & end), subcellular localisation, association with genetic disease (online Mendelian inheritance in man, OMIM), number of PubMed articles, evidence for NuRD interaction in PubMed and BioGRID, and cell line dependency from Human DepMap,.

**Supplementary Table 2.** List of bonds formed in the ZNF512B-RBBP4 3D structural model at the start of the molecular dynamics simulation run (0 ns). The amino acid residues in ZNF512B and RBBP4 involved in the interaction, as well as the distance, category and type of bond are described.

**Supplementary Table 3**. List of bonds formed in the ZNF512B-RBBP4 3D structural model at the end of the molecular dynamics simulation run (20 ns). The amino acid residues in ZNF512B and RBBP4 involved in the interaction, as well as the distance, category and type of bond are described.

**Supplementary Table 4.** List of oligonucleotides used in the study.

**Supplementary Table 5.** Lists of all differentially expressed genes (DEGs) detected after *ZNF512B* shRNA knockdown for 72 h in HEK293T cells. Each *ZNF512B* shRNA was compared to the non-targeting shRNA control.
