## Supplementary Table 4 for "The ZNF512B-NuRD complex co-ordinates a neural-specific gene expression program"

**Supplementary Table 4.** List of oligonucleotides used in this study

| **Name** | **Sequence (5’-3’)** | **Purpose** |
| --- | --- | --- |
| ZNF512B_shRNA1 _FWD | CCGGTGAGCAAACCTGTCACTATTGCTCG  AGCAATAGTGACAGGTTTGCTCATTTTTG | ZNF512B knockdown |
| ZNF512B_shRNA1 _REV | AATTCAAAAATGAGCAAACCTGTCACTAT  TGCTCGAGCAATAGTGACAGGTTTGCTCA | ZNF512B knockdown |
| ZNF512B_shRNA2 _FWD | CCGGCTATCACTTGTAACCATATATCTC  GAGATATATGGTTACAAGTGATAGTTTTTG | ZNF512B knockdown |
| ZNF512B_shRNA2_REV | AATTCAAAAACTATCACTTGTAACCATAT  ATCTCGAGATATATGGTTACAAGTGATAG | ZNF512B knockdown |
| qZNF512B_FWD | CACATGGAGGTGTGTCAGAAGC | qRT-PCR |
| qZNF512B_REV | GCCATAGTGTGGTAGTTGAGGC | qRT-PCR |
| qNANOG_FWD | CTCCAACATCCTGAACCTCAGC | qRT-PCR |
| qNANOG_REV | CGTCACACCATTGCTATTCTTCG | qRT-PCR |
| qPAX6_FWD | TCCATCAGTTCCAACGGAGAAG | qRT-PCR |
| qPAX6_REV | GTGGAATTGGTTGGTAGACACTG | qRT-PCR |
| qGAPDH_FWD | TGCACCACCAACTGCTTAGC | qRT-PCR |
| qGAPDH_REV | GGCATGGACTGTGGTCATGAG | qRT-PCR |
